# A three-dimensional imaging pipeline for the detection of vasculogenic mimicry in mouse tumor models

**DOI:** 10.64898/2026.10.07.756703

**Authors:** Lauren Deighton, Sarah M. Pearsall, Fatime Qosaj, Mike Mitchell, Griselda Awanis, IMAXT Cancer Grand Challenge Consortium, Dario Bressan, Caroline Dive, Kathryn L. Simpson, Gregory J. Hannon, Carlos Caldas, Kirsty Sawicka, Ian G. Cannell

## Abstract

Vasculogenic mimicry (VM) describes the ability for tumor cells to mimic endothelial cells and form tumor-lined blood vessels. Although VM is associated with poor prognosis, resistance to therapeutics and metastasis, its presence, relevance and functionality have been debated, largely due to the techniques employed to determine its presence. Dual CD31 and periodic acid-Schiff (PAS) histological staining, the method widely used to detect VM within tumor tissue, is hampered by issues with artifacts of 2D sections and does not assess active blood flow through vessels and hence cannot confirm vessel functionality. Here, we present a 3D imaging pipeline to detect functionally perfused VM and host vessels within mouse tumor models. Mice tumor models are intravenously injected with fluorescently-labelled tomato lectin which binds to the lumen of functional blood vessels. These tumors are thickly sliced, optically-cleared, stained with endothelial markers and imaged using light-sheet microscopy. We apply this imaging modality to image tumors from six breast cancer patient-derived tissue xenograft models, an orthotopic mammary cancer model and two small cell lung cancer circulating-tumor cell explant models, where VM vessels, mosaic vessels and host-VM vessel junctions can be detected. To our knowledge, this is the largest collection of 3D VM vessel images to date, providing convincing evidence of the presence and functionality of VM within tumors and defining the level of evidence necessary to detect VM *in vivo*.

## Introduction

To sustain the high metabolic demands of growing solid tumors, they must establish their own vascularization network (Hanahan & Weinberg, 2000). It was believed that the majority of these vessels were endothelial cell-lined and generated via angiogenesis. In angiogenesis and in response to hypoxia, pro-angiogenic signals such as vascular endothelial growth factor (VEGF) stimulate the branching of new vessels from pre-existing vessels (Saman et al., 2020). However, it has since emerged that tumor vessels can be formed by a multitude of alternative mechanisms, including vasculogenic mimicry (VM) (Ribatti & Pezzella, 2021). In VM, tumor cells acquire endothelial-like characteristics following hypoxia or therapeutic pressure and reorganise the extracellular matrix (ECM) to form tumor-lined vessels (Hendrix et al., 2003; Luo et al., 2020). These vessels are stabilized through cell-cell and cell-ECM adhesion molecules such as vascular endothelial (VE) cadherin and integrin β1 (Delgado-Bellido et al., 2023; Hendrix et al., 2001; Mingo et al., 2025; Pearsall et al., 2023; Williamson et al., 2016), and blood flow is maintained via the secretion of anti-coagulants (Ruf et al., 2003; Wagenblast et al., 2015). VM was first described in aggressive uveal melanoma, whereby the tumor cells could form tubular structures when grown on matrix *in vitro* and glycoprotein-rich vessels that stained positively for periodic acid-Schiff (PAS) reagent, devoid of endothelial cells, could be detected within tissue (Maniotis et al., 1999). Since then, VM has been reported in many cancer types where it has been associated with poor prognosis (Baeten et al., 2009; Cao et al., 2013; T. Liu et al., 2014; W. B. Liu et al., 2011; Maniotis et al., 1999; Sood et al., 2002; Wang et al., 2013; Williamson et al., 2016; Zhang et al., 2009), metastasis (Sun et al., 2006; Wagenblast et al., 2015) and therapeutic resistance (Cannell et al., 2023; Lezcano et al., 2014; Uceda-Castro et al., 2023; Williamson et al., 2016). Specifically, VM may serve as an alternative vascularization method when tumors are treated with anti-angiogenic therapy (AAT) and hence may contribute to the poor clinical translation of AAT (Coleman et al., 2017; Earl et al., 2017; Gilbert et al., 2014; Haibe et al., 2020; Salazar et al., 2020; Shepherd et al., 2022). Pre-clinical studies in breast and melanoma suggest that combined inhibition of VM and angiogenesis may be the best therapeutic strategy to halt the growth of tumors via blood vessel manipulation (Cannell et al., 2023; Delgado-Bellido et al., 2023). Despite this, the mechanism of VM *in vivo* remains poorly understood and consequently there are a lack of promising therapeutic targets to inhibit these deadly vessels (Fernández-Cortés et al., 2019).

*In vitro,* VM capability is typically assessed by the ability of tumor cells to form hollow, tubular structures when grown on ECM (Santander et al., 2026). Within tumor histological sections, it has become standard to define VM networks by PAS-positivity and the absence of an endothelial marker, such as CD31. However, this method has three main drawbacks which raise doubts about the presence, relevance and functionality of VM vessels within tumors (McDonald et al., 2000). Firstly, the field currently lacks a robust and specific biomarker that labels only the tumor cells lining VM channels and hence we rely on PAS staining. However, PAS not only stains glycoproteins present within the basement membrane of vessels, but also other carbohydrate structures within tumors such as mucin deposits, cell surface molecules and ECM (Aalto & Collan, 1986; Mcmanus, 1948). Hence, vessel-like, PAS-positive structures need to be detected which can be subjective and time-consuming. Secondly, definition of a tumor-lined vessel by the absence of one endothelial cell marker in a single 2D tissue section is technically challenging and does not prove endothelial cells or other host cells are not present across the entire vessel lumen. Lastly, dual CD31/PAS stains cannot prove that the identified VM vessels are actively perfused with blood from the host circulation and hence contribute to blood flow within the tumor. To overcome these limitations, we have previously injected intravenous perfusion agents, such as tomato lectin which binds to glycoproteins present in the lumen of vessels (Robertson et al., 2015), to detect VM networks in both 2D and 3D, using small cell lung cancer (SCLC) circulating-tumor-cell derived xenograft (CDX) models (Pearsall et al., 2023) and mouse breast cancer models (Cannell et al., 2023), respectively. To visualize VM networks in 3D, mice bearing VM-capable mammary cancer 4T1-T tumors were intravenously injected with fluorescently-labelled tomato lectin. 1mm-thick tumor slices were optically cleared, stained for CD31 and imaged using light-sheet microscopy (LSM). Lectin-positive, CD31-negative VM networks could be identified which accounted for around 30% of the total vessels within peri-necrotic regions (Cannell et al., 2023). Within the same study and using the same imaging method, fluorescently-labelled tumor cells could be visualized around a lectin-positive, CD31-negative VM vessel that connected to a CD31-positive vessel (Cannell et al., 2023). To date, we believe this is the most compelling evidence that VM vessels are tumor-lined and can permit blood flow.

Here, we aim to establish the most convincing *in vivo* VM detection modality and apply it to a series of different cancer models. Improved ability to detect VM in tissues and robustly identify VM-capable models could serve as platforms to better understand the mechanism of VM *in vivo* and hence aid in the development of therapeutics to target them.

## Results

### VM vessel detection using CD31/PAS stains overestimates actively perfused VM vessels

CD31/PAS staining, the current gold standard for detection of VM within tissue samples, was performed across six breast cancer patient-derived tissue xenograft (PDTX) models (Fig 1A). The models display heterogeneous PAS-positive structures. Models STG316 and STG321 have large PAS-positive islands which resemble ECM secreted from pericytes or fibroblasts (Fig 1A). Conversely, models HCI010 and STG201 have cell surface molecules which stain positively for PAS (Fig 1A). This heterogeneity and non-specificity of PAS staining renders it challenging to differentiate between VM vessels and other carbohydrate-rich structures within the tumors reproducibly between models. Additionally, from these stains alone, it is impossible to know if any suspected VM vessels are actively perfused with blood from the circulation. Despite this, and with the addition of an adjacent H&E stain, putative VM vessels which are PAS-positive, CD31-negative and contain red blood cells were detected within two of the models, AB630 and HCI010 (Fig 1B).

**Figure 1:**
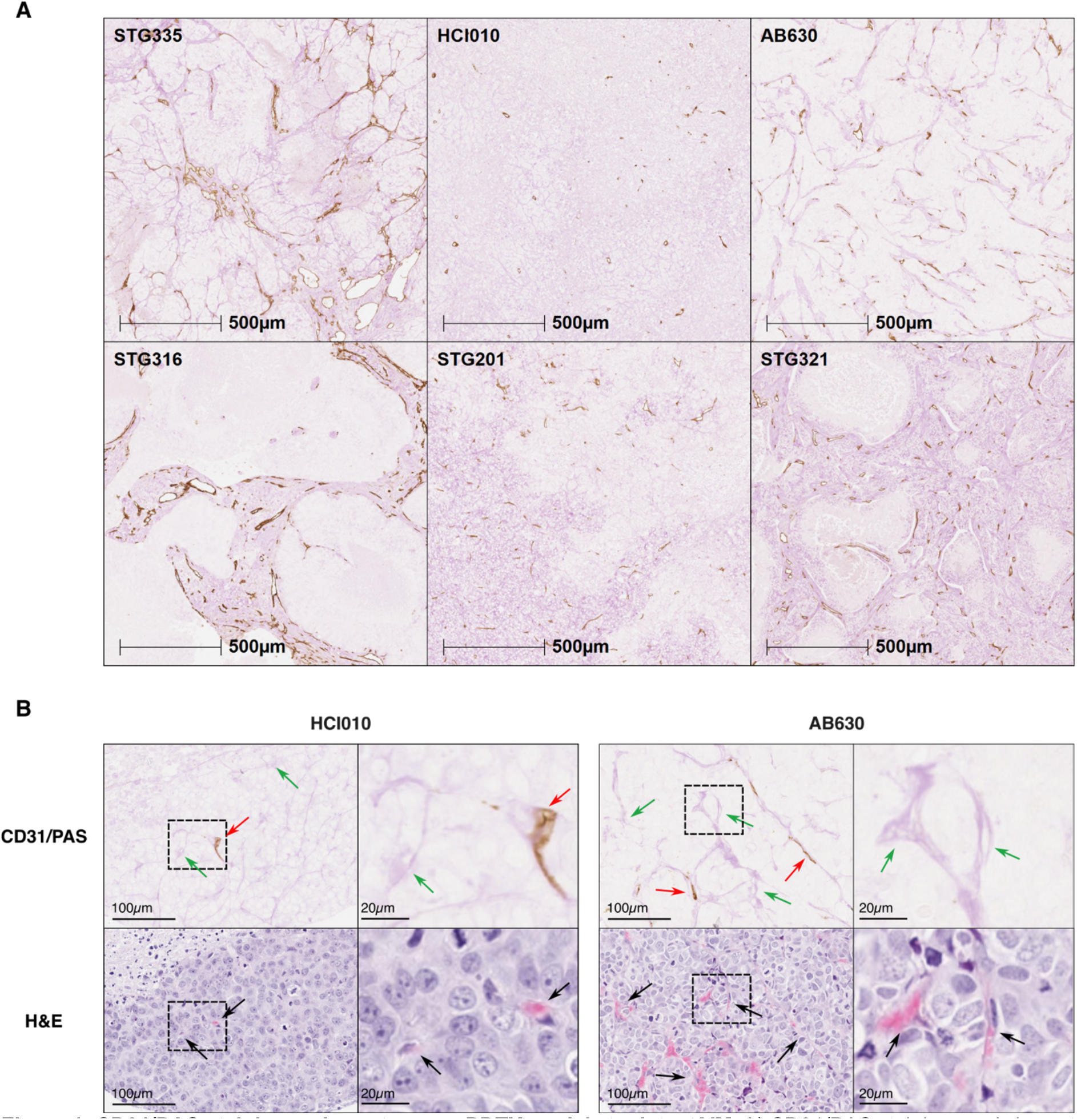
CD31/PAS staining on breast cancer PDTX models to detect VM. A) CD31/PAS staining on six breast cancer PDTX models. CD31-positive tissue (brown), PAS-positive tissue (pink). B) CD31/PAS staining (top) with adjacent H&E staining (bottom). CD31-negative, PAS-positive vessels (green arrows), that contain red blood cells (black arrows) can be detected amongst CD31-positive vessels (red arrows) in two PDTX models. Dotted square denotes a zoomed region.

To determine how many PAS-positive, CD31-negative, vessel-like structures are perfused with blood, mice bearing PDTX tumors were intravenously injected with fluorescently-labelled tomato lectin. Adjacent sections from the tumors were either stained with CD31/PAS; or CD31 alone and imaged for fluorescence to detect the lectin (Fig 2A). Lectin-positive, CD31-negative and PAS-positive vessels could be detected (Fig 2B); however, these were less abundant than quantified VM vessels from the CD31/PAS stains alone (Fig 2C). On average, around 16% of the PAS-positive, CD31-negative structures were lectin-positive and in model STG201 this was as low as 1.4% (Fig 2C). This suggests that VM detection using CD31/PAS stains significantly overestimates the number of functional VM vessels within tumor tissue sections, highlighting the need for better VM detection methods.

**Figure 2:**
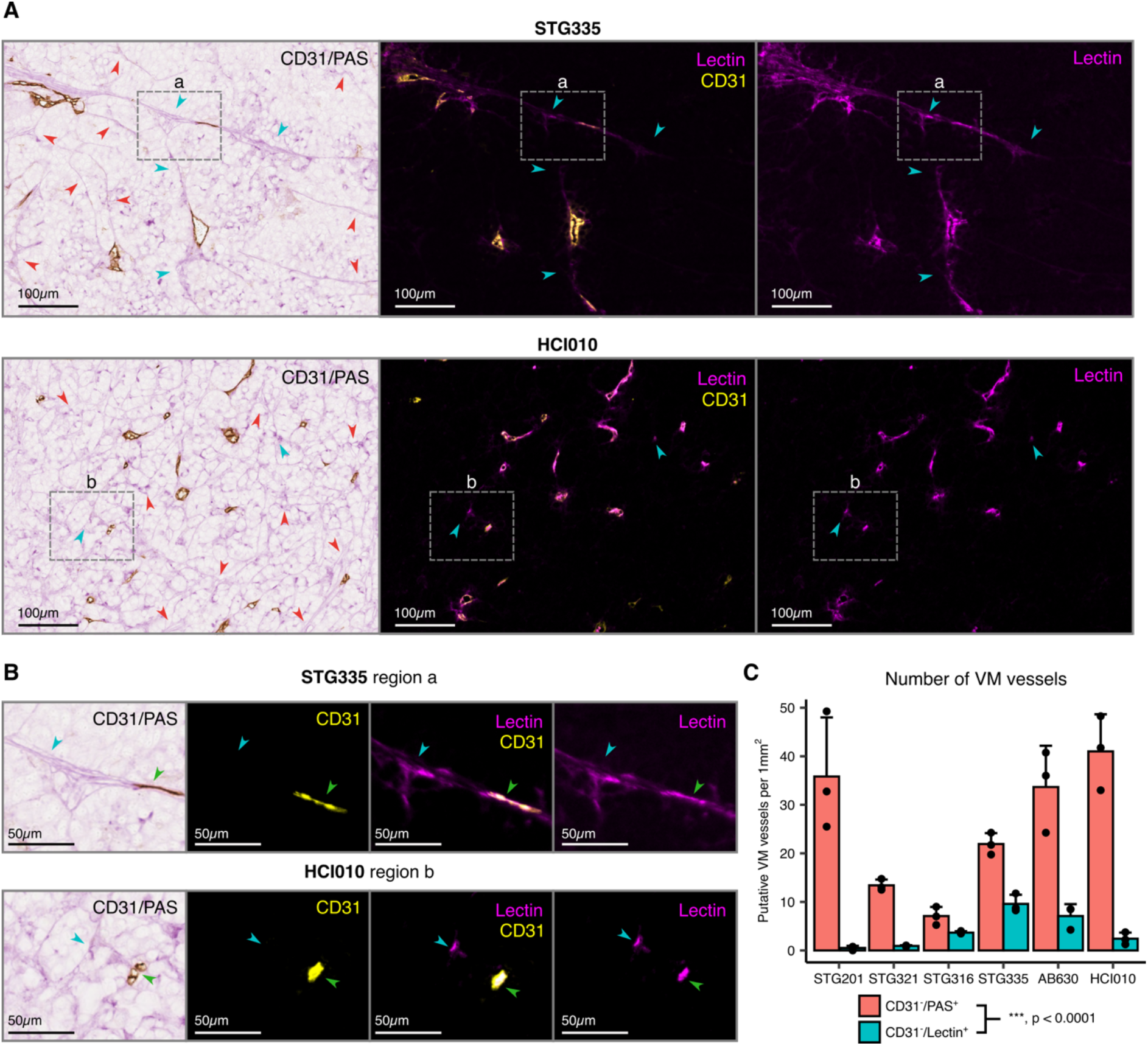
Comparison between CD31/PAS and CD31/lectin VM quantification in 2D images. A) Representative CD31/PAS (left panel) and CD31/lectin (middle and right panel) images from serial sections. Lectin (magenta)-perfused, CD31 (yellow)-negative, vessel-like structures (blue arrowheads) and CD31-negative, PAS-positive, vessel-like structures identified within CD31/PAS stains which are lectin-negative (red arrows). B) Zoomed regions a-b from A. Lectin-positive, PAS-positive, CD31-negative VM vessels (blue arrowheads) can be seen amongst lectin-positive, CD31-positive endothelial vessels (green arrowheads). C) The number of CD31-negative, PAS-positive, vessel-like structures identified per 1mm^2^ within CD31/PAS stains (red) and the number of CD31-negative, lectin-positive, vessel-like structures per 1mm^2^ within adjacent CD31/lectin images (blue) per model. n=3 representative areas from different tumors. A linear mixed-effects model was used to compare measurements obtained using CD31/PAS and CD31/lectin staining for VM quantification across the PDTX models (p = 2.5 × 10⁻⁸).

### VM vessels can be detected in breast cancer PDTX models in 3D

Tumors are known to have leaky vasculature and hence 2D visualization of perfused vessels cannot confirm CD31 negativity across the whole vessel diameter of putative VM vessels. Additionally, a single plane may not be representative of the whole tumor and does not allow vessel connectivity and structure to be observed. Therefore, we aimed to visualize these lectin-positive putative VM vessels in 3D. Six breast cancer PDTX models were intravenously injected with lectin conjugated to DyLight 649 in triplicate, then sliced to 1mm thickness, optically cleared, stained with anti-CD31 and imaged using LSM (Fig 3). Alternative fluorophores conjugated to lectin or perfusion agents such as fluorescent space-filling gels also enabled vessel detection, but lectin conjugated to DyLight 649 displayed the strongest signal to background ratio on all tumors imaged by LSM (Supp Fig 1).

**Figure 3:**
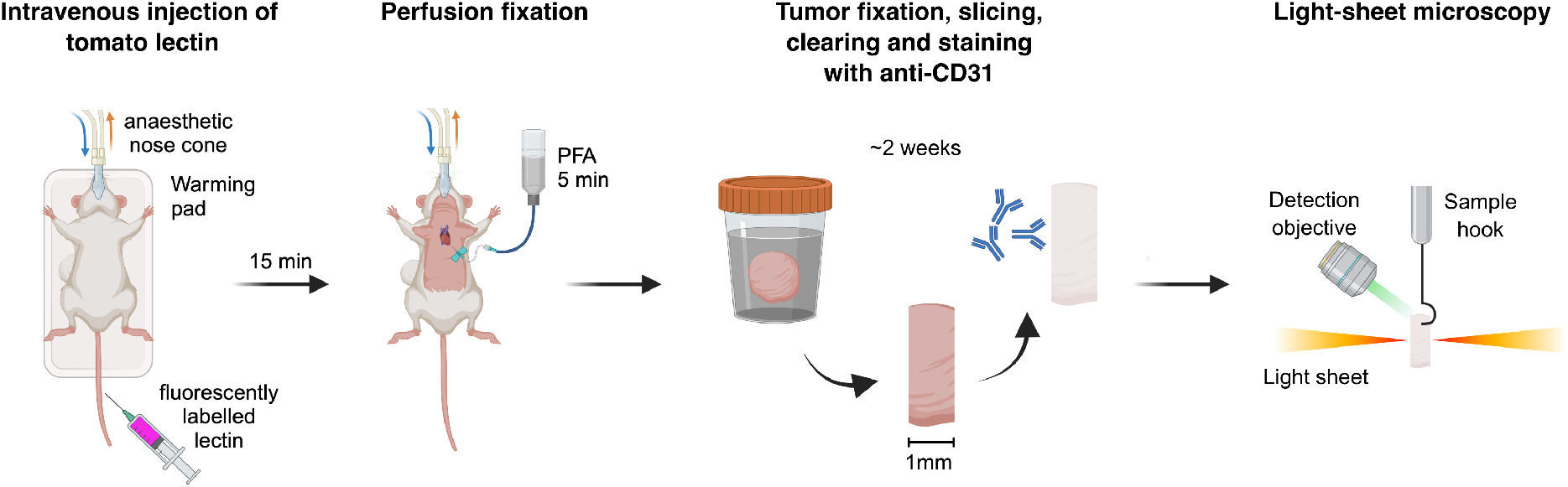
Overview of the 3D VM imaging pipeline. Under anaesthesia and on a heated mat, mice harbouring tumors are intravenously injected with fluorescently-labelled lectin. 15 minutes later, the mice are perfused with 4% paraformaldehyde (PFA) for 5 minutes and the tumors are collected and placed into 4% PFA for 24 hours. The tumors are sliced to 1mm thickness, optically cleared and stained with anti-CD31 before imaging using light-sheet microscopy.

Applying the 3D VM imaging pipeline to each PDTX model demonstrated that the vasculature was heterogeneously structured (Fig 4), and this heterogeneity was conserved over mouse replicates (Supp Fig 2). For example, the vessels in STG316 cluster together and HCI010 has smaller individual vessels that accumulate around the periphery of the tumor (Supp Fig 2). Four out of the six models consistently displayed lectin-positive, CD31-negative VM vessels (Fig 4, Supp Fig 3), with mosaic vessels (Supp Fig 3A), host-to-VM vessel junctions (Fig 4, Supp video 1) and blood pools observed (Supp Fig 3B, Supp video 1), which are structures that have been rarely documented, particularly in 3D. Cropped regions of these 3D tumor vasculature images can be segmented and assessed for CD31-positvity before being quantified (Supp Fig 3). The percentage of VM vessels varies between regions of the same tumor and between tumors (Supp Fig 3), similarly to what was observed in 4T1-T tumors previously, where VM vessel abundance increased near areas of necrosis (Cannell et al., 2023). Altogether, this demonstrates that we have established a 3D vessel imaging pipeline that can detect and quantify perfused VM vessels, alongside other vessel structures, such as mosaic vessels, in breast cancer PDTX models.

**Figure 4:**
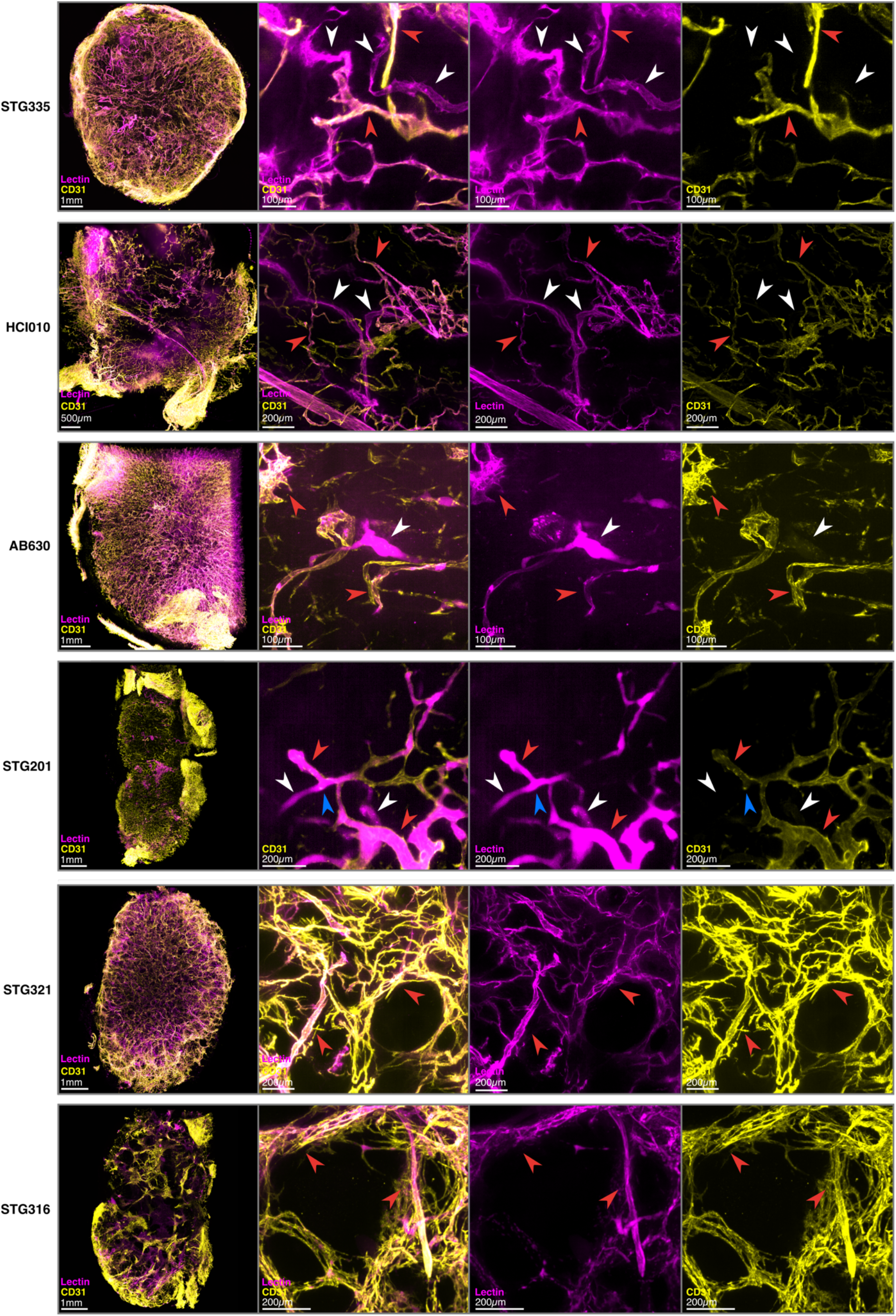
3D vasculature of breast cancer PDTX models. LSM 3D images (left) of a lectin perfused (magenta), CD31-stained (yellow) tumor slice from six different PDTX models. Cropped max intensity z-projections (right three panels) of lectin-positive, CD31-negative vessels (white arrows) and lectin-positive vessels with positive CD31 staining (red arrows) can be seen. A VM-to-host vessel junction can be observed within model STG201 (blue arrowhead).

### The 3D vessel imaging pipeline can be applied to other models

Three 4T1-T mouse mammary cancer tumors were processed and imaged using the 3D VM imaging pipeline (Fig 5). Lectin-positive, CD31-negative VM vessels could be detected within all three replicates (Fig 5C) and in keeping with previous findings, the abundance of these varied between tumors and within different regions of the same tumor (Fig 5B). Furthermore, VM vessels and mosaic vessels could be detected within two SCLC CDX models (Fig 6, Supp video 2), that were previously shown to contain VM vessels by CD31/PAS staining (Pearsall et al., 2023).

**Figure 5:**
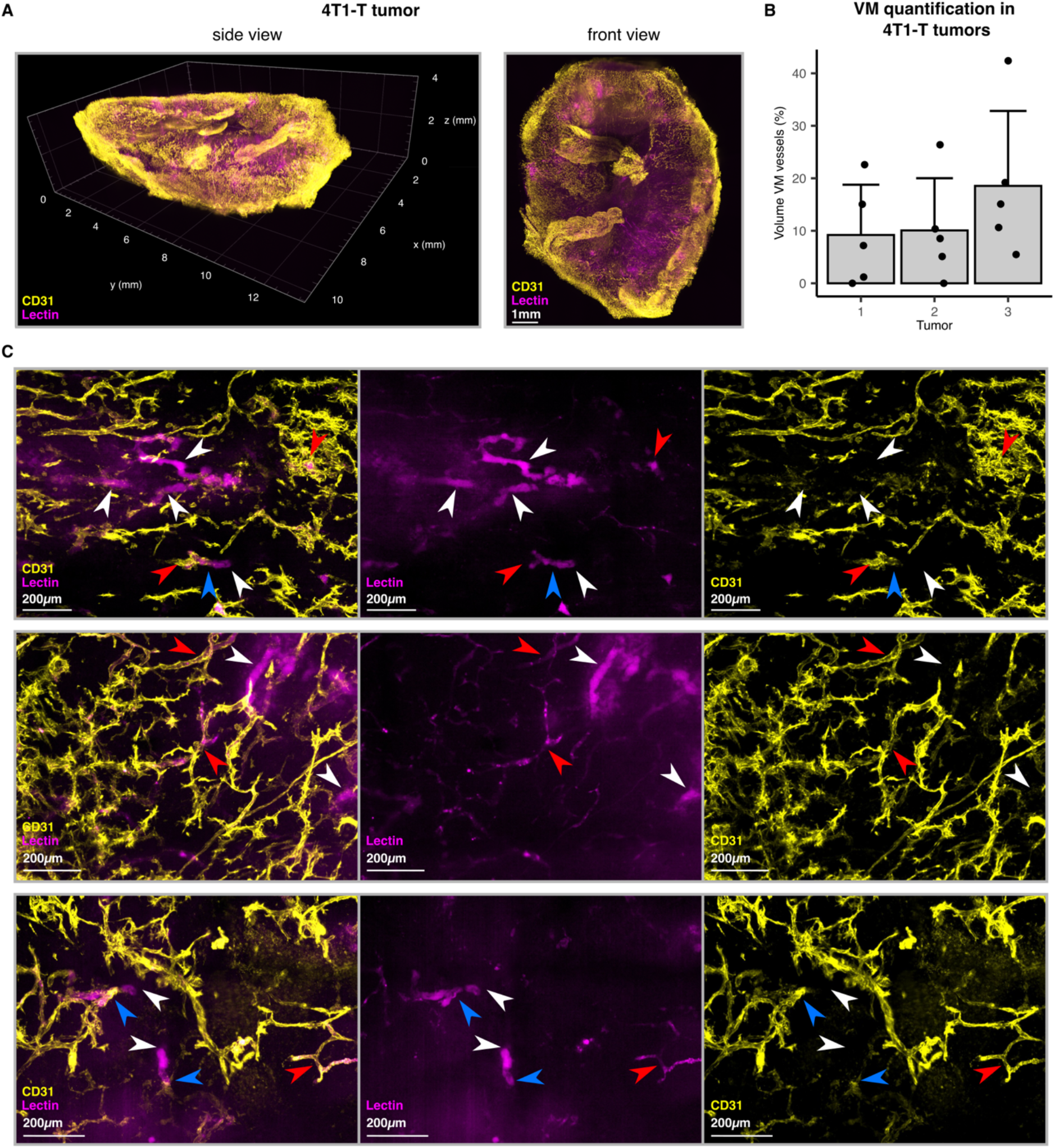
3D vessel imaging in 4T1-T tumors. A) Whole slice view of lectin-perfused, CD31-stained 4T1-T cleared tumor slices imaged by LSM. B) VM quantification of five regions from three separate 4T1-T tumors imaged by the 3D VM imaging pipeline on LSM. Error bars = sd. C) Max projections of cropped regions of lectin-perfused, CD31-stained 4T1-T cleared tumor slices imaged by LSM. Lectin-positive, CD31-negative VM vessels (white arrowheads) can be detected amongst lectin-positive, CD31-postive host vessels (red arrowheads). Some VM-to-host vessel junctions can be observed (blue arrowheads).

**Figure 6:**
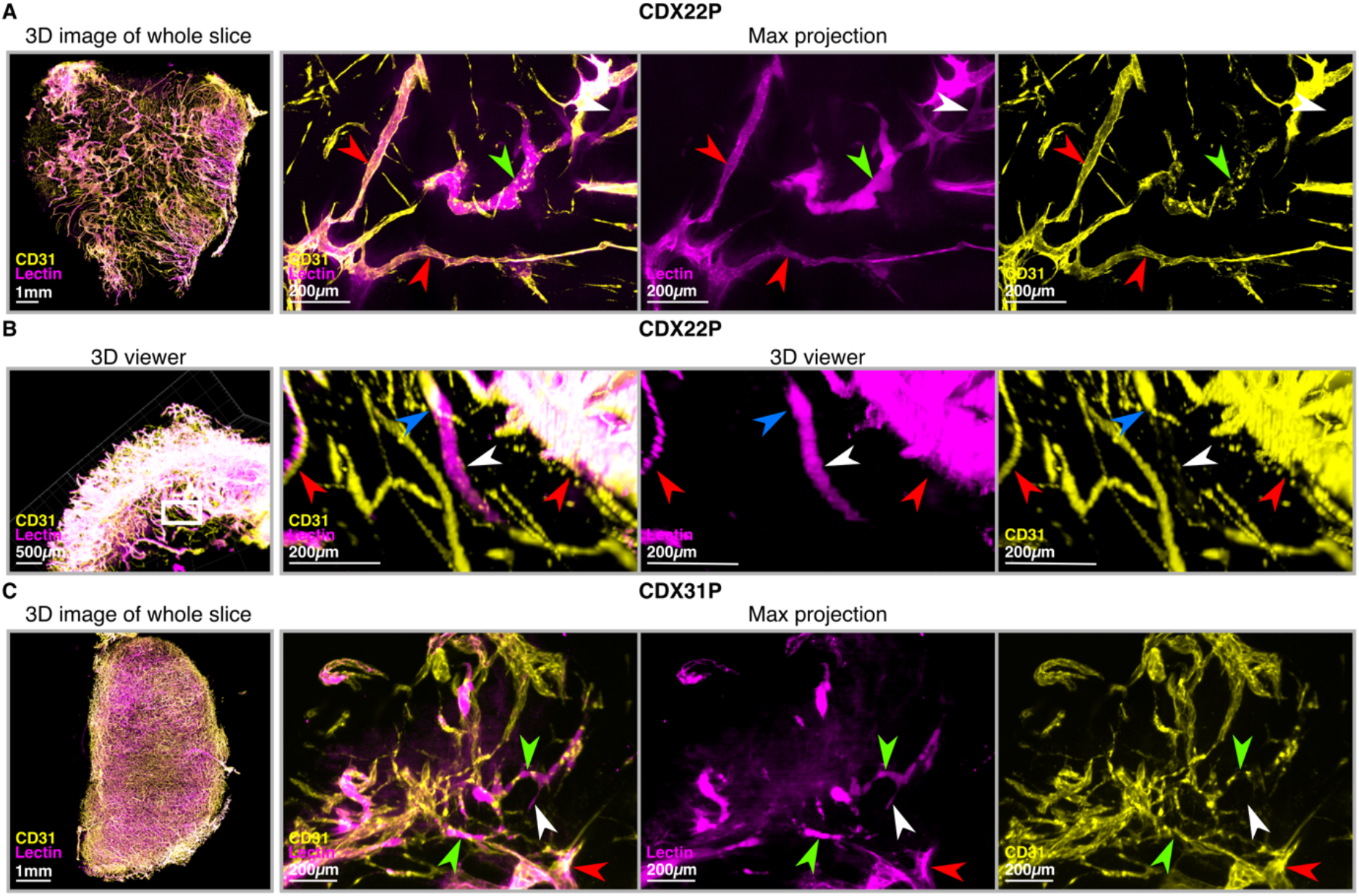
3D vessel imaging in SCLC CDX models. A) Lectin-perfused, CD31-stained 3D imaging of an optically cleared slice of a CDX22P SCLC CDX tumor (left). Max projection of a cropped region of the same tumor displaying VM (white arrowheads), mosaic (green arrowheads) and host (red arrowheads) vessels (top right). B) 3D view of a lectin-positive, CD31-negative VM vessel (white arrowhead) connected to a CD31-positive host vessel (blue arrowhead), surrounded by perfused host vessels (red arrowheads) in CDX22P. C) Lectin-perfused, CD31-stained 3D imaging of an optically cleared slice of a CDX31P SCLC CDX tumor (left). Max projection of a cropped region of the same tumor displaying VM (white arrowheads), mosaic (green arrowheads) and host (red arrowheads) vessels (right).

### VM vessels have no mouse cells surrounding them in breast cancer PDTX model HCI010

CD31 negative staining alone may not be sufficient to prove that perfused vessels are lined with tumor cells; and are not lined by CD31-negative endothelial cells or other host cells capable of forming tubules, such as monocytes (Anghelina et al., 2006; Barnett et al., 2016). Previously, we circumvented this by using fluorescently labelled tumor cells to display tumor cell-positive, lectin-positive, CD31-negative VM vessels (Cannell et al., 2023). However, in models where survival is low and genetic modification is challenging *in vitro*, such as in PDTXs, fluorescent labelling of tumor cells is not feasible. Instead, we chose to label the mouse cells by utilising the NSG-EGFP transgenic mouse where all mouse cells are labelled with EGFP. We implanted VM-capable breast cancer PDTX model HCI010 into an NSG-EGFP mouse, intravenously injected fluorescently-labelled lectin and imaged vessels using our 3D LSM pipeline (Fig 7, Supp video 3). In this model, we can identify lectin-positive, CD31-negative VM vessels that are also EGFP-negative suggesting that no mouse cells surround them and that they are tumor-lined (Fig 7).

**Figure 7:**
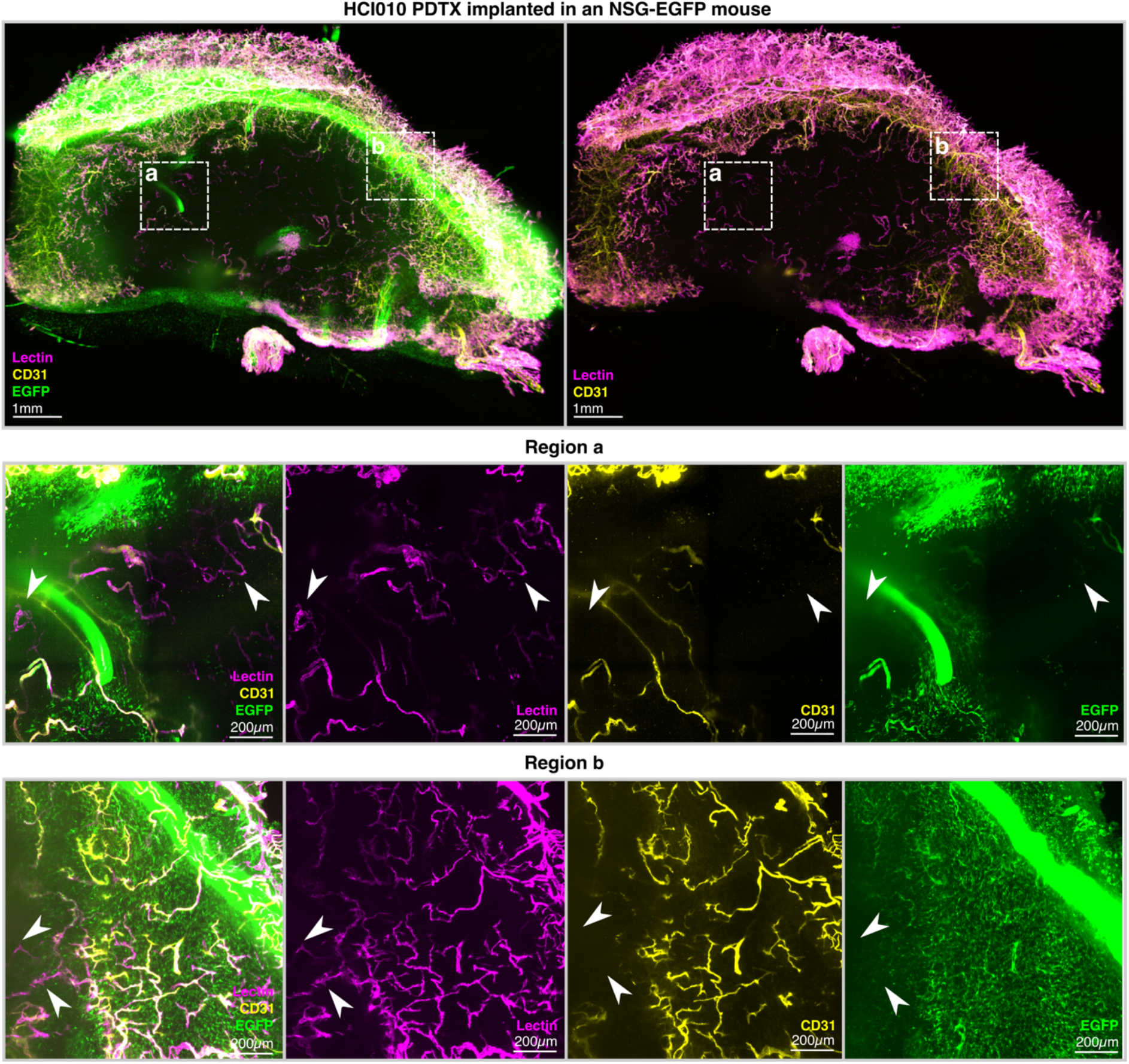
VM vessels within HCI010 implanted into an NSG-EGFP mouse. LSM 3D images (top) of a lectin-perfused (magenta), CD31-stained (yellow) HCI010 tumor slice implanted in an NSG-EGFP mouse (green). Max intensity z-projections (bottom) of 500μm of region a and 100μm of region b. Lectin-positive, CD31-negative, mouse-cell negative VM vessels can be detected (white arrows).

## Discussion

Since the discovery of VM in uveal melanoma in 1999 (Maniotis et al., 1999), it has been associated with poor prognosis (Cao et al., 2013), therapeutic resistance (Cannell et al., 2023; Lezcano et al., 2014; Uceda-Castro et al., 2023; Williamson et al., 2016) and metastasis (Sun et al., 2006; Wagenblast et al., 2015) in a variety of cancer types. Although a few key molecules have been identified mainly through *in vitro* assays (Luo et al., 2020; Seftor et al., 2012), the mechanism remains unclear and hence potential therapeutic strategies to target VM vessels are limited. Additionally, technical challenges of detecting VM vessels in tumors have limited our understanding of the mechanism *in vivo* and attracted scepticism in the field (McDonald et al., 2000).

### Methods to pinpoint VM vessels *in vivo* and their limitations

Here, we assessed current methods to pinpoint VM vessels in tumor tissues across six breast cancer PDTX models and summarised the advantages and disadvantages in Table 1. As predicted, the widely used CD31/PAS stains overestimate the abundance of functional VM vessels (Fig. 2C). It is unknown whether this overestimation is due to the non-specific nature of PAS which leads to the incorrect identification of PAS-positive ‘vessel-like’ structures as VM vessels; or if these vessel-like endothelial-deficient PAS structures are actually tubules that were not perfused during the timeframe of the experiment. It is plausible that they could be tumor-lined tubules which have not connected to host vasculature yet, and hence VM vessel precursors. Additionally, the PDTX models studied display heterogeneity in the PAS-positive structures (Fig. 1A), making it difficult to be consistent with what is defined as ‘vessel-like’ between models. However, AI that is trained using adjacent lectin-perfused images, may be used to predict perfused PAS-positive VM structures from CD31/PAS stains. There is currently no robust and specific biomarker for identification of the tumor cells lining VM vessels. Therefore, despite the technical limitations, CD31/PAS staining on 2D sections remains the best approach for detecting VM vessels in tumors where perfusion agent injection is not feasible, such as in human samples. CD31/PAS stains are inexpensive and can be performed easily on FFPE tissue routinely collected in preclinical studies or clinical trials. Furthermore, CD31-negative, PAS-positive, vessel-like structures are associated with poor prognosis across multiple cancers (Cao et al., 2013) and hence have prognostic value, even if the structures identified may not be functional tumor-lined vessels.

**Table 1:** Comparison of methods to visualize VM vessels *in vivo*.

| Method | Pros | Cons |
| --- | --- | --- |
| **2D**<br>**CD31/PAS** | • Inexpensive.<br>• Can be performed on human samples and routinely collected FFPE samples. | • PAS stain is non-specific and variable between samples.<br>• Laborious and subjective to detect and quantify VM.<br>• Cannot confirm vessels are perfused with blood from the circulation. |
| **2D**<br>**CD31/lectin** | • Confirms blood perfusion status of vessels.<br>• Easier and less subjective to detect and quantify than CD31/PAS stains<br>• Best option for combining with spatial transcriptomics or highly multiplexed immunostaining | • 2D section cannot determine if the vessel is CD31 negative across the full cross-section.<br>• Can only perform on animal models.<br>• Unknown if lectin binds to all VM vessels. |
| **3D**<br>**CD31/lectin** | • Confirms blood perfusion status of vessels and can visualize connectivity to host vessels.<br>• CD31 negativity is confirmed over the whole vessel diameter. | • Can only perform on animal models.<br>• Unknown if lectin binds to all VM vessels.<br>• CD31 antibody needs to penetrate 3D sample.<br>• Clearing, imaging and quantification is expensive and slow. |

The use of perfusion agents such as tomato lectin greatly increased confidence of vessel functionality and allowed for the visualization of VM vessels in both 2D (Fig. 2) and 3D (Figs. 4-7) across mouse models and human patient-derived models. Tomato lectin has been widely used to visualize endothelial-lined blood vessels across various tissues, where it binds to glycoproteins present within the lumen of these vessels (Bergers et al., 2003; Lokmic & Mitchell, 2011; Robertson et al., 2015). Lectin-positive, CD31-negative vessels could be detected within this study suggesting that tomato lectin is also capable of binding to VM vessels. However, it is unknown whether all VM vessels display the glycoproteins that lectin binds to and hence VM vessel detection using this method may be underestimated. Additionally, since lectin binds to the lumen of vessels, segmentation of larger vessels for quantification can be difficult. Fluorescently labelled, space-filling gels would be an alternative perfusion agent that would display no binding bias and may be easier to segment. Gels such as polymer resin PU4ii have labelled perfused vasculature clearly within the mouse brain using confocal microscopy (Krucker et al., 2006). Using fluorophore SeTau665 dispersed within PU4ii resin and our clearing and imaging set up, we could also clearly detect vessels within the mouse brain (data not shown). However, in a tumor from the same mouse as the imaged brain, the signal was not strong enough to overcome the autofluorescence of the tumor (Supp Fig 1). Alternative clearing, fluorophores or imaging systems may be needed to further optimise this to act as an alternative to tomato lectin.

A limitation of all these methods to visualize VM is the reliance on the lack of CD31 staining to detect the absence of endothelial cells. In 2D, absence of CD31 staining in a single plane cannot confirm that the whole lumen of the vessel is negative. Furthermore, in 3D, efficiency of staining is a technical challenge since the antibody must penetrate the dense tumor slice and hence tumor areas must be carefully selected for strong CD31 staining prior to investigation. The limitations are also biological since endothelial vessels can have varying expression of CD31 dependant on the tissue location and vessel type (Pusztaszeri et al., 2005; Trimm & Red-Horse, 2022) and non-endothelial cells such as monocytes have been observed to form vessel-like structures that are perfused with blood in tissues (Anghelina et al., 2006; Barnett et al., 2016). Such vessels would stain negatively for CD31 and be falsely characterised as VM using this method. Here, we addressed this limitation with the use of the NSG-EGFP transgenic mouse, whereby lectin-positive, CD31-negative vessels that were also EGFP negative could be identified, confirming the absence of mouse cells around the suspected VM vessels.

### Future perspective

Here, we present a pipeline for imaging the tumor vasculature in 3D and show that we can optically clear and stain a variety of different models, including patient-derived xenografts. Although applied here to detect VM vessels, alternative antibodies, fluorescently labelled tumor cells, or transgenic mice could be used to investigate a wide range of other questions in 3D.

Using our 3D vessel imaging modality, we detected mosaic vessels and VM-host junctions, structures which have rarely been documented in 3D. Although the VM vessels must connect to host vasculature to be perfused, the exact mechanism of this and the molecular connections required between the cells is not known. Traditionally, it has been proposed that the VM-capable tumor cells reorganise the ECM to generate their own tubules, however the complete redundancy of endothelial cells to support this process has not been confirmed or documented *in vivo*. The prospect that VM vessels become tumor-lined by displacement or death of endothelial cells with host-derived vessels is a theory which has not been explored. In support of this, breast cancer cells have been visualized to integrate into established endothelial-lined tubules using *in vitro* 3D microvessel models (Silvestri et al., 2020). Further studying the interactions between tumor and endothelial cells to form functional VM vessels and understanding how to inhibit them may be critical in preventing the tumor bypass inefficiencies in the host vasculature system.

A lack of understanding of the VM mechanism is currently hindering our therapeutic strategies to inhibit them. The method we present here confidently identifies which models are VM-capable. Hence, combined with further characterization such as utilising single-cell or spatial transcriptomics, VM-capable models could be interrogated to identify key drivers of VM or VM-specific biomarkers. Here, the imaging pipeline is used across a series of different models to generate the largest showcase of functional VM vessel examples in 3D to date. Used further, this imaging tool could help us understand more about these fascinating but deadly vessels and how to target them therapeutically.

## Methods and Materials

### Resource Availability

#### Lead contact

Further information and requests for resources and reagents should be directed to and will be fulfilled by the lead contact, Ian Cannell.

#### Materials availability

Resources and reagents requests should be directed to, and will be fulfilled by, the lead contact.

#### Data and code availability

- Original data will be shared by the lead contact upon request.
- This paper does not report original code.
- Any additional information required to reanalyze the data reported in this paper is available from the lead contact upon request.

### Tumor xenograft formation

All mouse work was conducted according to the UK Home Office regulations (Project license #PAD85403A and #PP4872079) and under the CRUK Cambridge Institute establishment license approved by the Animal Welfare and Ethical Review Board.

#### Breast cancer PDTX tumors

PDTX models were established and passaged as in (Bruna et al., 2016). Briefly, fragments sized around ∼3mm^3^ were defrosted rapidly in a 37°C water bath, washed in Dulbecco’s Modified Eagle Medium (DMEM) (Thermo Fisher #41965039), placed into Matrigel (Corning #356231) and kept on ice until implantation. Female NSG (Charles River #614) or NOD.Cg-Prkdcscid Il2rgtm1Wjl Tg (CAG-EGFP) 1Osb/SzJ (NSG-EGFP, The Jackson Laboratory #021937) mice typically aged ∼6 weeks were anaesthetised with isoflurane and tumor fragments were inserted subcutaneously into the flank.

#### 4T1-T tumors

4T1-T cell line generated in-house (Wagenblast et al., 2015) with a Rosa26 knock-out, was cultured in DMEM (Thermo Fisher #41965039) supplemented with 10% HI FBS (Thermo Fisher #A5256801) in a humidified incubator at 37°C and 5% CO2. 50μl of a 1:1 mixture containing 50,000 cells in PBS and Matrigel (Corning #356231) was injected into the fourth mammary fat pad of female BALB/C mice (Charles River #028) aged ∼8 weeks.

#### SCLC CDX tumors

SCLC CDX models were established and passaged as described in (Simpson et al., 2020). Briefly, 100,000 dissociated tumor cells in 100µl 1:1 RPMI:Matrigel were injected subcutaneously into the flank of female NSG (Charles River #614) mice aged ∼8 weeks.

### Histology of tumor tissue

#### Formalin-fixed paraffin embedded (FFPE) tissue

Collected tumors were incubated for 24 hours in 10% neutral buffered formalin (NBF, Fisher Scientific #13191184) at room temperature with rotation. Tumors were washed twice for 2 hours in PBS with rotation and then placed in 70% ethanol. Tumors were dehydrated and embedded within paraffin using the following programme: 70% ethanol (30 mins x 1), 96% ethanol (30 mins x 2), 100% (30 mins x 2), xylene (30 mins x 3) and paraffin wax (1 hour, 60°C). Tumors were then placed into moulds in paraffin wax and cooled on a cold plate.

#### FFPE tissue sectioning, slide preparation and staining

3μm sections of chosen FFPE blocks were sectioned and baked at 60°C for one hour. Slides were dewaxed and rehydrated through washes of 100% xylene (10 mins x 2), 100% ethanol (5 mins x 2) and 70% ethanol (5 mins x 1) and then placed into water for subsequent processing.

#### Haematoxylin and Eosin (H&E) staining

Incubations were performed automatically using a Leica Multistainer ST5020 using the following programme: Harris haematoxylin (2 mins), water (2 mins), 2% acid alcohol (20 seconds), water (5 mins), eosin (7 mins), water (15 seconds), 50% ethanol with 0.01% tween (20 seconds), 70% ethanol with 0.01% tween (20 seconds), 100% ethanol (30 seconds x 2) and 100% Xylene (5 mins x 2). Slides were coverslipped and imaged at 20X resolution using a Lecia Aperio AT2 whole slide scanner.

#### CD31/PAS staining

Immunohistochemistry (IHC) staining of CD31 (Cell Signalling #77699, 1:100 dilution) was performed using a Leica Bond autostainer with 20 minutes antigen retrieval at 100°C in Tris-EDTA retrieval buffer (Leica #RE7119) and a Bond Polymer Refine Detection System (Leica Biosystems #DS9800). Following IHC staining, slides were incubated in the following solutions: ultrapure water (20 seconds), 0.5% periodic acid (Sigma Aldrich #P0430)(5 mins), ultrapure water (20 seconds x 3), Schiff’s reagent (Fisher Scientific #J/7300/PB08) (30 mins) and ultrapure water (20 seconds x 3). Slides were coverslipped and imaged at 20X resolution using a Leica Aperio AT2 whole slide scanner. CD31-negative, PAS-positive, vessel-like structures were manually detected from representative 2mm x 2mm areas for each sample. A vessel-like structure was defined as either a long stretch of PAS until a new connection or a PAS-positive lumen.

#### CD31 immunofluorescence

Slides were incubated for 10 minutes at 100°C in tri-sodium citrate buffer (10mM tri-sodium citrate in water, pH 6.0) in the microwave followed by a brief wash in PBS and incubation in 0.2% Triton X-100 in PBS for 30 minutes. Slides were washed in PBS for 5 minutes 3 times with gentle rocking. Blocking buffer (10% Goat Serum (Merck #S26-LITER), 150mM Maleimide (Merck #129585), 100mM Ammonium Chloride in PBS) was added and slides incubated for 2 hours at room temperature. Blocking buffer was removed and CD31 antibody (Cell Signalling #77699) diluted 1:100 in staining buffer (5% Goat Serum, 100mM Ammonium Chloride in PBS) was added. Slides were incubated overnight at 4°C within a humidity chamber and the following day washed 3 times in PBS for 5 minutes. 1μg/ml DAPI (BD Biosciences #564907) and fluorophore-conjugated secondary antibody (anti-rabbit Alexa Fluor 555: ThermoFisher #A21428) diluted 1:300 in staining buffer was added to the slides and incubated at room temperature in the dark for 2 hours. Slides were washed in PBS 3 times for 5 minutes with gentle rocking. Slides were coverslipped using ProLong diamond antifade mountant (ThermoFisher Scientific #P36965) and cured at room temperature for 24 hours before imaging at 20X resolution using the Zeiss Axioscan Z1 whole slide scanner. Using QuPath and a pixel intensity thresholder, lectin-positive objects larger than 30μm^2^ were segmented and CD31-positivity determined to more easily detect vessels in an unbiased manner. On a representative 2mm x 2mm area of tissue, it was manually determined which CD31-negative, lectin-positive objects were distinct vessel-like structures

### 3D tumor vessel imaging

#### Mouse vessel labelling

Mice bearing tumors sized between 500-1000mm^3^ were incubated at 37°C for 10 minutes before anaesthetic induction with isoflurane in a heated chamber. Once anaesthetised, mice were transferred to a heated mat within a fume cupboard and remained under anaesthetic via a nose cone. Breathing rate was carefully monitored and once at a steady rate, 150μl of Lycopersicon Esculentum (Tomato) lectin conjugated to DyLight 649 (2BScientific #DL-1178-1) was injected into the tail vein. Alternative fluorophores DyLight 488 (Thermo Scientific #L32470), DyLight 594 (Thermo Scientific #L32471), Texas Red (Thermo Scientific #L32482) were also tested during pipeline optimization. 15 minutes later, the mouse’s rib cage was opened to allow for injection of a needle into the left ventricle and an excision was made to the right atrium. 4% PFA (Alfa Aesar #043368.9M) diluted in PBS was pumped at a rate of 1ml/minute using a peristaltic pump for 5 minutes.

#### Alternative vessel labelling agents

##### CM-Dil

100mg of CellTracker CM-DiI (ThermoFisher Scientific #C7000) was resuspended in 16.7ml of 100% ethanol to create the CM-Dil stock solution. A dilutant solution was made by diluting mixing a 5% (w/v) glucose solution in water with PBS in a 1:4 ratio. The mouse was anaesthetised with isoflurane and placed in a fume cupboard and remained under anaesthetic via a nose cone. PBS with 32mg/L heparin was pumped into the left ventricle using a peristaltic pump at a rate of 1ml/minute for 5 minutes. A CM-DiI working solution was prepared immediately before use by adding 200μl of CM-DiI stock solution to 10ml dilutant solution and pumped at a rate of 1ml/minute for 5 minutes. 4% PFA (Alfa Aesar #043368.9M) diluted in PBS was pumped at a rate of 1ml/minute using a peristaltic pump for 5 minutes.

##### Resin containing fluorescent dye or beads

Mice were anaesthetised with isoflurane and PBS with 32mg/L heparin was pumped into the left ventricle using a peristaltic at a rate of 1ml/minute for 5 minutes, followed by 4% PFA (Alfa Aesar #043368.9M) diluted in PBS at a rate of 1ml/minute pump for 5 minutes. Using a syringe, either an acrylamide/bead gel mix or PU4ii resin containing SeTau 665 dye was prepared as below and injected slowly into the left ventricle. Acrylamide/bead gel: A 10ml acrylamide/bead mix containing 2.5ml 40% acrylamide/bis solution (Bio-Rad #1610144), 6.6μl of N,N,N′,N′-Tetramethylethylenediamine (TEMED) and 200μl of 2μM QTracker quantum dots 655 (ThermoFisher Scientific #Q21021MP) or 0.2μm yellow/green fluorospheres (ThermoFisher Scientific #F8811) was prepared. Immediately before injection, 66μl 10% ammonium persulfate (APS) solution was added to the acrylamide/bead solution. PU4ii resin with SeTau 665: A resin/dye mix containing 2.5g polyurethane resin (PU4ii, VasQTec), 3g 2-butanone and 300μl 1mg/ml SeTau 665 dye (SETA BioMedicals #K9-4119) dissolved in dimethylformamide (DMF) was prepared and immediately before injection, 0.95g of harder (VasQTec) was added.

#### Tumor sample preparation, clearing and staining

After mouse sacrifice and tumor collection, tumors were incubated in 4% paraformaldehyde (PFA, Alfa Aesar #043368.9M) in PBS at 4°C with rotation for 24 hours. Tumors were washed three times with PBS for 2 hours at 4°C with rotation before being sliced to 1mm thickness using a tissue slicer (Zivic Instruments #HSRS005-1). Tumor slices were incubated in 50% CUBIC-L (TCI Chemicals #T3740) for 6 hours at 37°C with shaking at 100 rpm. Slices were transferred to 100% CUBIC-L and incubated at 37°C with shaking at 100 rpm for a further 4-7 days with changing the solution every 1-2 days. If required, samples would also be incubated with 100% CUBIC-1 solution (25% (wt) urea, 25% (wt) Quadrol, 15% (wt) Triton X-100 in distilled water) for a further 4-7 days with solution changes every 1-2 days to fully clear the sample. The tumor slice was then washed three times with PBS for 2 hours with rotation. Slices were incubated at room temperature with gentle rocking with CD31 antibody (Cell Signalling #77699) diluted 1:100 in PBS for 4 days, with exchange of antibody solution and flipping of the tumor slice after 2 days. The slice was washed three times in PBS containing 0.1% Tween for 2 hours with rocking and incubated with anti-rabbit goat secondary antibody conjugated to Alexa Fluor 555 (ThermoFisher Scientific #A-21429) diluted 1:300 in PBS and incubated for 3 days at room temperature with gentle rocking. Slices were washed three times in PBS containing 0.1% Tween for 2 hours with rocking, then incubated in 50% CUBIC-R+ (TCI Chemicals #T3741) at room temperature with rotation for 6 hours, followed by incubation with rotation at room temperature in 100% CUBIC-R+. All the incubation steps described above were performed with light protection around the samples to reduce fluorophore quenching.

#### Sample imaging, processing and visualization

Cleared samples were imaged immersed within CUBIC-R+ (TCI Chemicals #T3741) using a 5X dry objective lens (EC Plan-NEOFLUAR 5x/0.16 ZEISS) and sCMOS PCO edge cameras within the Zeiss Lightsheet Z.1 system (Carl Zeiss, Germany) with 10% overlap between tiles, 10μm z-steps and dual side illumination with pivot scan. For imaging of DyLight 488, yellow/green fluorospheres, GFP or EGFP, a 488 nm excitation laser was used, split using a SBS LP 490 splitter and detected through a BP 505-545 filter. For imaging of Alexa Fluor 555, Texas Red, DyLight 594 or CM-DiI, a 561 nm excitation laser was used, split using a SBS LP 640 splitter and detected through a BP 575-615 filter. For imaging of DyLight 649, QTracker 655 dots or SeTau 665, a 638 nm excitation laser was used, split using a SBS LP 640 splitter and detected through a LP 660 filter. Laser intensity and exposure time were adjusted dependent on the signal, but remained the same for sample cohorts quantified together. Tiles were stitched and visualized within the Arivis Vision 4D software. For max projection images, regions of interest were cropped and the z-planes flattened within FIJI by extraction of the max signal intensity.

#### VM quantification from LSM images

3-5 regions from each imaged tumor slice were cropped. Each region measured 2mm x 2mm and contained 50 z-stacks (equating to 500μm depth) from planes which had strong CD31 staining. The sample names were randomised to reduce bias and reassigned after quantification. Using Arivis Vision 4D software, the lectin signal was denoised using gaussian blur set to a radius of 5μm and segmented using “Otsu thresholding for big structures”, based on extracting objects with the top 5% signal intensity within the whole image. Objects larger than 100μm^2^ were selected. The segmentation mask was used to calculate the total volume of perfused vasculature within each image and segments were manually determined whether they also have CD31 signal and regrouped accordingly. The volume of the lectin-positive, CD31-negative objects was used to calculate the percentage volume of VM vessels in respect to the total lectin-positive area.

## Acknowledgements

We would like to thank all members of the Hannon, Caldas and IMAXT labs at the CRUK Cambridge Institute (CI, RRID:SCR_028157), in particular Clare Rebbeck for assistance with home office animal licensing. Furthermore, we would like to thank all the core facilities at the CRUK CI, in particular; Histopathology and In Situ Hybridisation (RRID:SCR_028183) for preparation of FFPE blocks and performing histological sectioning, staining and imaging; Pre-Clinical Genome Editing (RRID:SCR_028661) for the implantation of PDTX models; the Microscopy core (RRID:SCR_028019), in particular Fadwa Joud, for training and maintenance of the light-sheet microscope; and members of the Biological Resources Unit (RRID:SCR_028671) for animal husbandry and care, specifically Mike Mitchell, for performing intravenous lectin injections and intra-cardiac perfusions. We are also thankful to all members of the Pre-Clinical Genome Editing core facility (RRID:SCR_028661), Caldas group and Dive group involved in the generation, characterization and biobanking of the PDTX and CDX models respectively.

This work was supported in part by the IMAXT Cancer Grand Challenges award (C9545/A24042 to G.J.H.) and a Cancer Research UK core award (A21143 to G.J.H.). This work was also supported in part by the Assistant Secretary of Defense Health Agency endorsed by the Department of Defense, through the FY17 Breast Cancer Research Program; Breakthrough Award (W81XWH1810703 to G.J.H. and I.G.C). Opinions, interpretations, conclusions, and recommendations are those of the authors and are not necessarily endorsed by the Department of Defense.

We are very grateful for the generosity of all the patients that donated samples for PDTX and CDX implantation. We are also thankful to all the staff at the Cambridge Breast Cancer Research Unit, CRUK Cambridge Cancer Centre, Cambridge University Hospital NHS Foundation Trust (funded by Addenbrooke’s Charitable Trust, Cambridge University Hospitals, and CRUK) and the NIHR Manchester Clinical Research Facility at The Christie Hospital for consenting patients and facilitating the timely collection of samples.

## Author Contributions

Conceptualization: L.D., C.C., D.B., K.S., I.G.C. and G.J.H. Data curation: L.D. Formal analysis: L.D. Funding acquisition: I.G.C., D.B., and G.J.H. Investigation: L.D., S.P., G.A., F.Q., and M.M. Methodology: L.D., S.P., I.G.C., F.Q., D.B., M.M., and K.S. Project administration: L.D., I.G.C., and K.S. Resources: G.A., K.L.S., C.D., C.C., and G.J.H. Software: L.D. Supervision: I.G.C., K.S., C.D., C.C., and G.J.H. Visualization: L.D. Writing - original draft: L.D. Writing - reviewing and editing: L.D., I.G.C., S.P., G.A., and K.S.

