## Supplementary Information for "A three-dimensional imaging pipeline for the detection of vasculogenic mimicry in mouse tumor models"

### Supplement

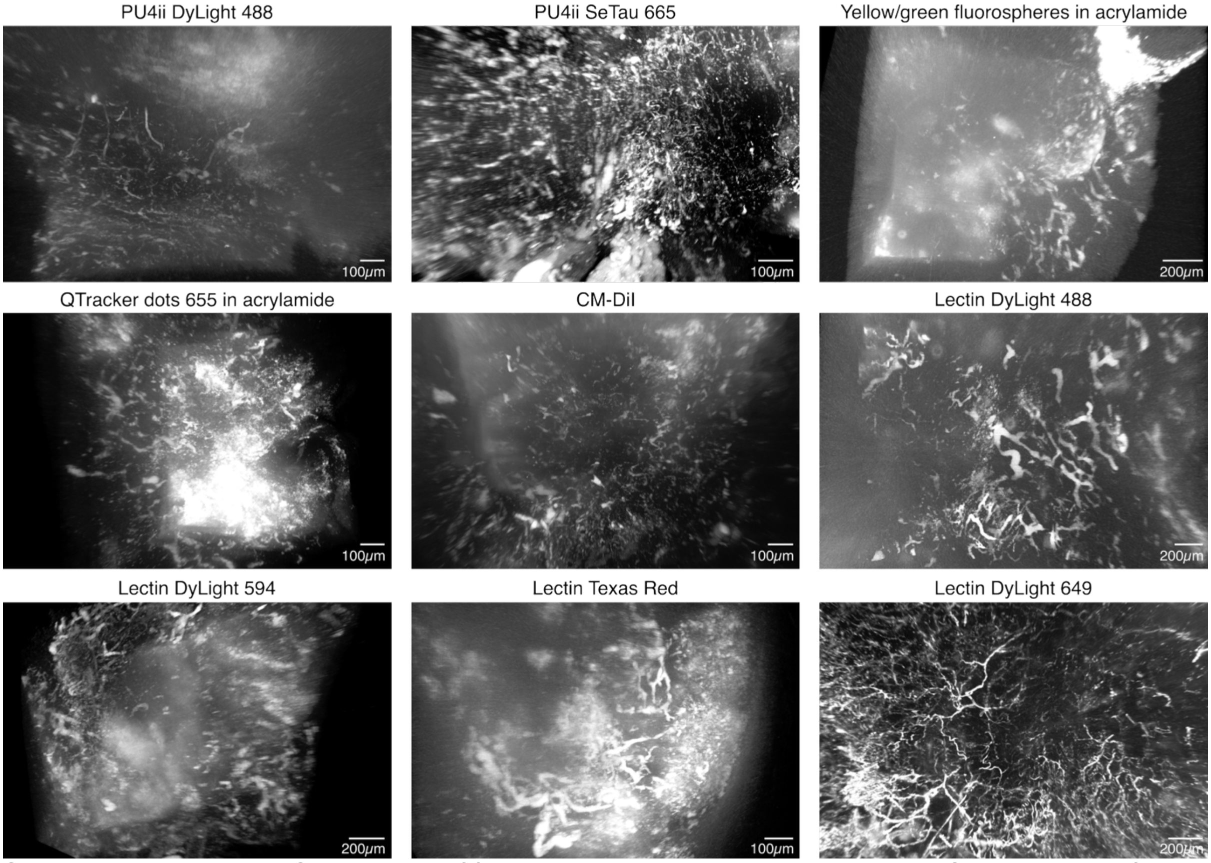


**Supplementary Figure 1: Optimization of functional vessel labelling agents using LSM.** 3D images of 4T1-T tumors perfused with different vessel labelling agents and imaged using LSM. PU4ii and acrylamide are space-filling fluorescently labelled gels and the others are fluorescently-labelled vessel-binding agents.

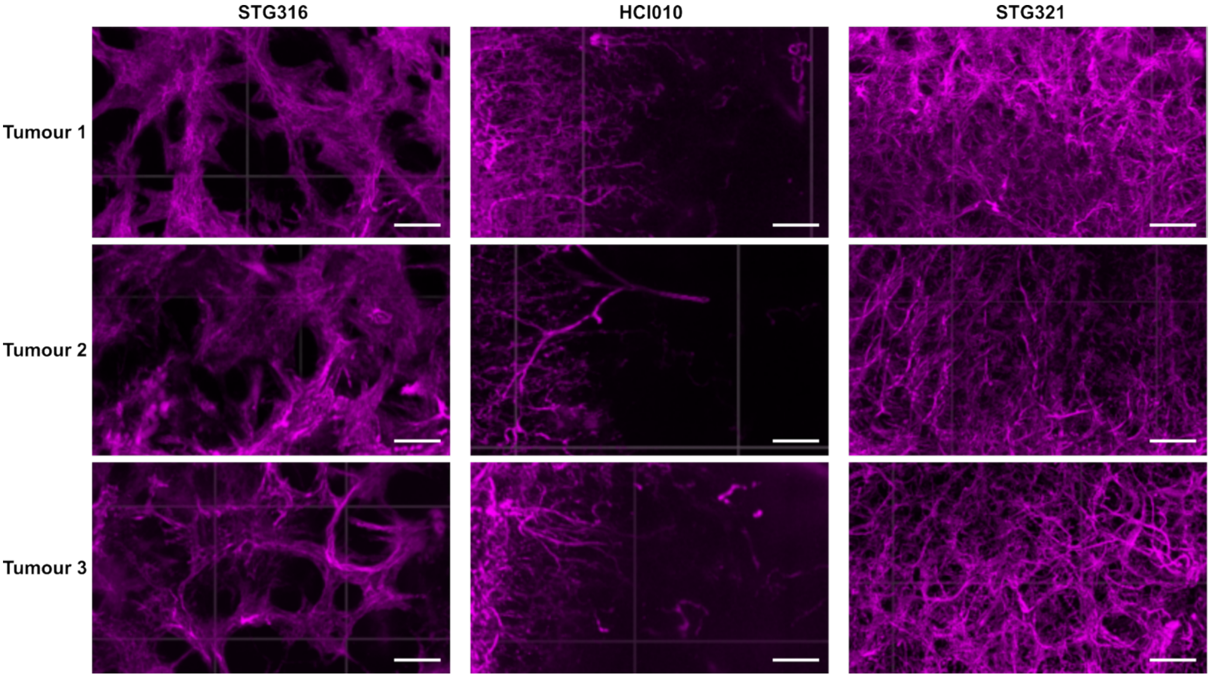

**Supplementary Figure 2: PDTX models display vasculature heterogeneity between models which is conserved over replicates**. 3D LSM images of three replicates of three different PDTX models intravenously injected with lectin (magenta). Scale bars = 100µm.

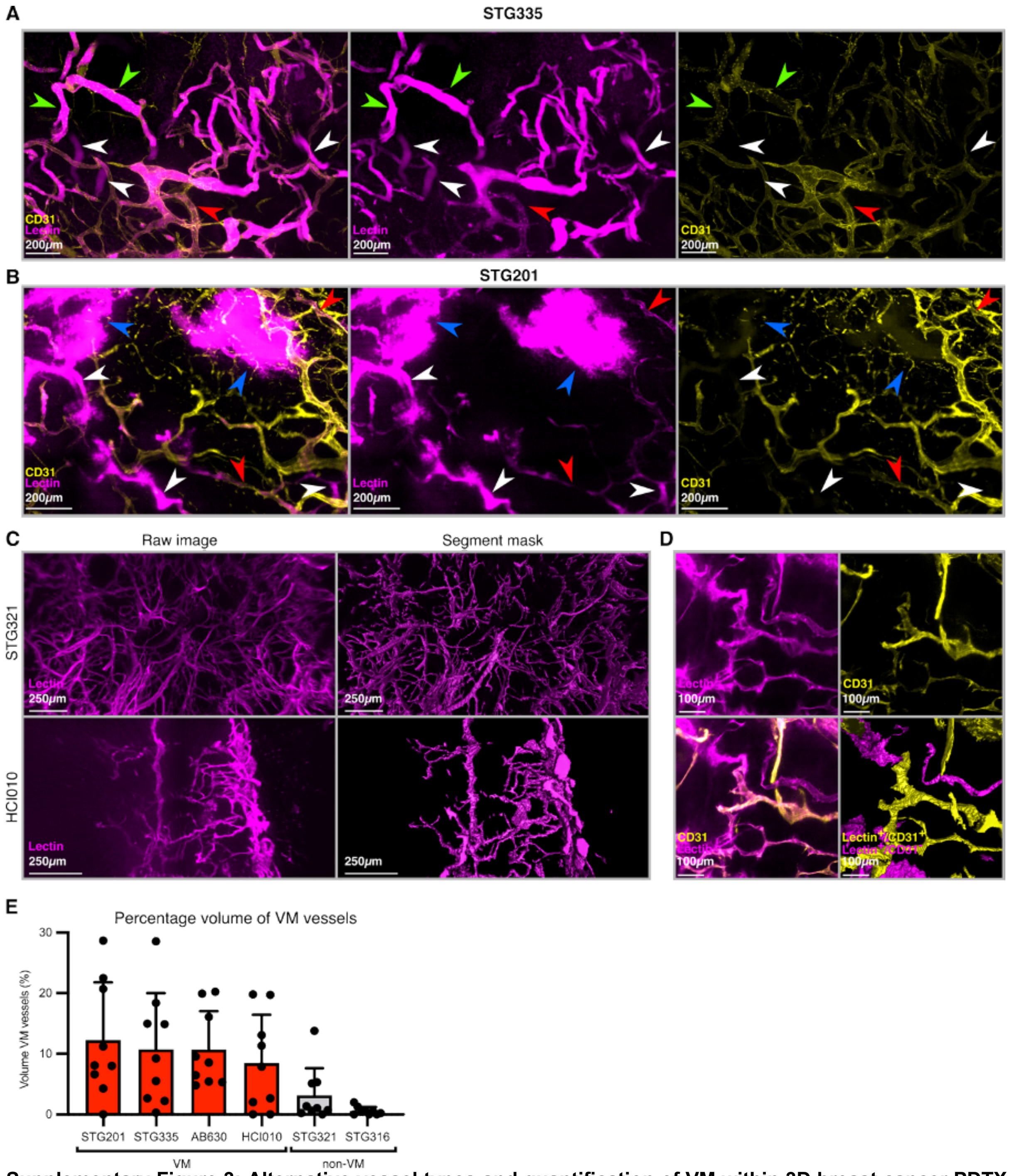


**Supplementary Figure 3: Alternative vessel types and quantification of VM within 3D breast cancer PDTX models.** A) Max projection of a cropped region of STG335 imaged using the 3D VM imaging pipeline. Lectin-positive, CD31-negative VM vessels (white arrowheads), perfused host vessels (red arrowhead) and perfused mosaic vessels (green arrow heads) can be detected. B) Max projection of a cropped region of STG201 imaged using the 3D VM imaging pipeline. Lectin-positive, CD31-negative VM vessels (white arrowheads), perfused host vessels (red arrowhead) and large blood pools (blue arrowheads) can be detected. C) Representative examples of the segmentation of the lectin signal in two PDTX models, STG321 and HCI010. D) Example from STG335 of the raw images and the segmented and classified vessels (bottom right). Lectin-positive, CD31-negative VM vessels are displayed in magenta and lectin-positive, CD31-positive host vessels are displayed in yellow. E) Quantification of the percentage volume of VM vessels out of the total lectin-positive vessels in six breast cancer PDTX models imaged using the 3D VM imaging pipeline. Quantification was performed on three different tumors for each model with three regions cropped to 2mm x 2mm and 500μm depth for each tumor. Error bars = sd.

**IMAXT Cancer Grand Challenge Consortium Authors**

The following authors were part of the IMAXT Cancer Grand Challenge Consortium:

Bruno Albuquerque (1), Martina Alini (1), Heather Ashmore (1), Thomas Ashmore (1), Giorgia Battistoni (1), Dario Bressan (1), Ian Gordon Cannell (1), Hannah Casbolt (1), Lauren Deighton (1), Ilaria Falciatori (1), Carla Boquetale (1), Nikki Coutts (1), Chee Ying Sia (1), Atefeh Fatemi (1), Nicole Hemmer (1), Kui Hua (1), Cristina Jauset (1), Tatjana Kovačević (1), Claire M Mulvey (1), Natasha Narayanan (1), Fiona Nugent (1), Clare Rebbeck (1), Marta Paez Ribes (1), Isabella Pearsall (1), Sarah Pearsall (1), Fatime Qosaj (1), Kirsty Sawicka (1), Sophia A Wild (1), Elena Williams (1), Hamid Raza Ali (1), Samuel Aparicio (1), Emma Laks (2), Yangguang Li (2), Ciara H O'Flanagan (2), Austin Smith (2), Teresa Ruiz (2), Daniel Lai (2), Andrew Roth (2), Vinci Au (2), Caroline Baril (2), Sean Beatty (2), Shankar Balasubramanian (2), João CF Nogueira (1), Max Lee (1), Bernd Bodenmiller (4), Alina Bollhagen (4), Marcel Burger (4), Laura Kuett (4), Jonas Windhager (4), Edward S Boyden (4), Debarati Ghosh (5), Anubhav Sinha (5), Brett Pryor (5), Ruihan Zhang (5), Jack Lovell (5), Chi Zhang (5), Yangning Lu (5), Carlos Caldas (5), Alejandra Bruna (1), Maurizio Callari (1), Wendy Greenwood (1), Giulia Lerda (1), Yaniv Eyal Lubling (1), Oscar M Rueda (1), Abigail Shea (1), Owen Harris (1), Robby Becker (6), Natalie Duncan (6), Flaminia Grimaldi (6), Suvi Harris (6), Sara Lisa Vogl (6), Joanna Weselak (6), Johanna A Joyce (6), Spencer S Watson (7), John Marioni (7), Sohrab P Shah (1), Andrew McPherson (2), Ignacio Vázquez-García (2), Simon Tavaré (8), Khanh N Dinh (1), Russell Kunes (9), Nicholas A Walton (9), Mohammad Al Sa'd (10), Nick Chornay (10), Ali Dariush (10), Eduardo A González Solares (10), Carlos González-Fernández (10), Melis Irfan (10), Aybüke Küpcü Yoldaş (10), Alireza Molaeinezhad (10), Neil Millar (10), Leigh Smith (10), Tristan Whitmarsh (10), Xiaowei Zhuang (10), Jean Fan (11), Hsuan Lee (11), Leonardo A Sepúlveda (11), Chenglong Xia (11), Pu Zheng (11)

(1) Cancer Research UK Cambridge Institute, Li Ka Shing Centre, University of Cambridge, Cambridge CB2 0RE, UK
(2) Department of Molecular Oncology, BC Cancer, part of the Provincial Health Services Authority, Vancouver, BC, Canada
(3) Department of Chemistry, University of Cambridge, Lensfield Road, Cambridge, CB2 1EW, UK
(4) Department of Quantitative Biomedicine, University of Zurich, Zurich 8057, Switzerland
(5) McGovern Institute, Departments of Biological Engineering and Brainand Cognitive Sciences, Massachusetts Institute of Technology, Cambridge, Massachusetts, USA, and HHMI, Cambridge, Massachusetts, USA
(6) Súil Interactive Ltd, Dame Lane, Dublin, UK
(7) Department of Oncology and Ludwig Institute for Cancer Research, University of Lausanne, Lausanne, Switzerland
(8) Computational Oncology, Department of Epidemiology and Biostatistics, Memorial Sloan Kettering Cancer Center, New York, USA
(9) Herbert and Florence Irving Institute for Cancer Dynamics, Columbia University, New York, NY, USA
(10) Institute of Astronomy, University of Cambridge, Madingley Road, Cambridge, CB3 0HA, UK
(11) Howard Hughes Medical Institute, Harvard University, Cambridge, MA 02138, USA; Department of Physics, Harvard University, Cambridge, MA 02138, USA; Department of Chemistry and Chemical Biology, Harvard University, Cambridge, MA 02138, USA
